# Analysis of AlphaFold and molecular dynamics structure predictions of mutations in serpins

**DOI:** 10.1101/2023.01.31.526415

**Authors:** Pedro Garrido-Rodríguez, Miguel Carmena-Bargueño, María Eugenia de la Morena-Barrio, Carlos Bravo-Pérez, Belén de la Morena-Barrio, Rosa Cifuentes-Riquelme, María Luisa Lozano, Horacio Pérez-Sánchez, Javier Corral

## Abstract

Serine protease inhibitors (serpins) include thousands of structurally conserved proteins playing key roles in many organisms. Mutations affecting serpins may disturb their conformation, leading to inactive forms. Unfortunately, conformational consequences of mutations affecting serpins are difficult to predict. In this study we integrate experimental data of patients with mutations affecting one serpin with the predictions obtained by AlphaFold and molecular dynamics. Five *SERPINC1* mutations causing antithrombin deficiency, the strongest congenital thrombophilia, (p.Arg79Cys; p.Pro112Ser, p.Met283Val; p.Pro352insValPheLeuPro, and p.Glu241_Leu242delinsValLeuValLeuValAsnThrArgThr-Ser) were selected from a cohort of 350 unrelated patients based on functional, biochemical, and crystallographic evidence supporting a folding defect. AlphaFold gave an accurate prediction for the wild-type antithrombin structure. However, it only produced native structures for all variants, regardless of its complexity and *in-vivo* conformational consequences. Similarly, molecular dynamics of up to 1000 ns at temperatures that caused conformational transitions did not show significant changes in the native structure of wild-type or variants. In conclusion, one predictive tool of protein folding, AlphaFold, and a simulation method for analyzing the physical movements of atoms and molecules, molecular dynamics, force predictions into the native stressed conformation at conditions with experimental evidence supporting a conformational change to relaxed structures. It is necessary to improve predictive strategies for serpins that consider the conformational sensitivity of these molecules.

## Introduction

Serpins are a superfamily of proteins sharing a common, strongly conserved structural configuration required to inhibit serin proteases. Even serpins with no inhibitory activity share a single common core domain consisting of 3 *β* strands, 7-9 *α* helices, and a reactive center loop (RCL) for the interaction with target proteases [1]. Serpins are folded into a stressed, metastable native conformation with 5 *β* strands in the central A sheet. This configuration allows serpins to change their structure to a hyperstable form with 6 *β* strands in response to stimulus, usually, the cleavage of the RCL by their target protease [2]. This efficient suicidal mechanism explains the key role of serpins in a number of crucial systems based on proteolytic cascades in a wide range of species. It is the case of coagulation, inflammation or the complement system. Such structural sensitivity also makes serpins particularly vulnerable to even minor modifications caused by environmental conditions or missense mutations, which may cause a conformational instability with pathogenic consequences [3], as hyperstable conformations (latent, when the RCL is inserted into the own molecule, and polymers, if the insertion involves another molecule) with no inhibitory activity. Moreover, polymers may be toxic if accumulated in the cell [4]. These aberrant conformations, caused by a wide range of mutations affecting *SERPINA1*, *SERPINC1*, *SERPINI1* or *SERPING1*, are responsible of disorders of relevance, such as emphysema, thrombosis, neuropathies or angioedema, respectively [5]. Moreover, environmental conditions may also induce conformational changes in serpins [6].

One of the best-known serpins is antithrombin, probably the most important endogenous anticoagulant [7]. Its deficiency dramatically increases the risk of thrombosis, even when it impacts a single allele, acting as a dominant disorder. Antithrombin deficiency can be classified in two groups. Type I when the mutation, by different mechanisms, severely affects the levels of variant antithrombin in plasma. Type II, if the mutation does not severely impair the secretion of the antithrombin variant which has impaired or null anticoagulant activity. More than 386 different variants affecting *SERPINC1*, the gene encoding antithrombin, have been described as causative of antithrombin deficiency [8].

One great challenge for antithrombin and all other serpins is how to predict the consequences of any new mutation, aiming to identify those with a conformational effect [9]. Overall, prediction of 3D protein structures based on their sequences is a long-lasting problem in biochemistry [10]. Likewise, it has also been a noteworthy challenge for artificial intelligence (AI) systems since the advent of bioinformatics, and we can see nowadays that AI methods have started to slightly increase the accuracy of structural predictions [11]. In 2021, Jumper et al. [12] released AlphaFold, their AI model for protein structure prediction. The model developed by DeepMind has created high expectations in the community [13] because of its outstanding results on the 13th edition of Critical Assessment of Structure Prediction (CASP) [14] using the first version of the model [15], and more recently in CASP 14 [16] with the new AlphaFold 2 [12]. The model, now released to the community, is expected to ease the efforts needed to solve the 3D structures of several proteins yet unknown [17]. Nonetheless, the structural resolution of non-novel, mutated proteins could mean a huge step forward in several areas. Although its authors state it is not validated to predict mutational effects [18] and flipping an amino acid in the target protein to model a mutation will not work in AlphaFold [19, 20], it would be interesting to challenge the model to solve the conformational consequences of mutations affecting serpins, as these mutations cause a relevant structural change compared with the native conformation.

Molecular Dynamics (MD) simulation is an established and routinary methodology for the prediction of the dynamical evolution of biomolecular structure with atomic detail [21]. Indeed, first MD simulation of a simple protein system was carried out in 1977 [22], while recently MD allowed the simulation of the SARS-CoV-2 spike protein, implying millions of atoms [23]. Nonetheless, it should be mentioned that main limitations of the MD technique are related, among others, with the required computing time, which depends mainly on the system size and the trajectory length needed. Advances in hardware in the last decade allow nowadays to perform millisecond simulations for proteins of average size such as p53 [24]. Reaching longer timescales (i.e., seconds) is still a challenge since time dependent processes cannot be parallelized.

Thus, in this study we have explored AlphaFold’s and molecular dynamics predictions for different environmental conditions inducing conformational changes in antithrombin, as well as for natural mutants of antithrombin identified in our cohort of patients. All selected cases have variant antithrombins detected in plasma, ensuring both folding and secretion of variants, and were deeply studied using different biochemical methods providing information of their conformation, including a case whose crystal structure was determined.

## Materials and Methods

### Patients

For 23 years (1998-2021) our group has recruited one of the largest cohorts of unrelated patients with antithrombin deficiency (N = 350). After molecular analysis of *SERPINC1*, the gene encoding this key anticoagulant, 135 different gene defects were found in 250 patients. Experimental characterization of plasma antithrombin in these cases, described extensively elsewhere [25, 26, 27] included functional analysis of anti-FXa and anti-FIIa activities by using chromogenic methods, quantification of antigen levels by immunological methods, identification of forms with low heparin affinity by crossed immunoelectrophoresis in the presence of heparin, and separation of plasma antithrombins by native (in the presence and absence of 6M urea) and denaturing PAGE and western blotting, procedures that allow detection of aberrant forms of antithrombin and a semi-quantitative determination of the latent conformation [28].

For this study, we selected 5 different mutations leading to antithrombin variants detected in the plasma of carriers (Table 1). Indeed, we have experimental data of all variants, which may include the recombinant expression in eukaryotic cells of the mutated variant [29], its purification from plasma and in one case a crystallographic characterization. Moreover, this selection also aimed to cover the range of different mutations, from simple missense to the insertion of different number of residues in the structure of antithrombin.

**Table 1.** Functional antigenic and biochemical data of *SERPINC1* mutations identified in patients with antithrombin deficiency and selected for predictions using AlphaFold.

| Mutation | N | Anti-FXa | Antigen | Low<br>Hep<br>Aff<br>CIE | Latent<br>Native<br>Urea | Dimers | Other<br>data |
| --- | --- | --- | --- | --- | --- | --- | --- |
| *p.Arg79Cys* | 19 | 42.6 ± 9.8 | 91.6 ± 11.1 | Yes | - | - | - |
| *p.Pro112Ser* | 1 | 50 | 55 | - | - | Yes | - |
| *p.Met283Val* | 1 | 76 | 98 | Yes | Yes | - | - |
| *p.Pro352ins*<br>*ValPheLeuPro* | 5 | 47.5 ± 8.9 | 58.0 ± 1.7 | - | - | Yes | Verified by<br>proteomics |
| *p.Glu241-Leu242*<br>*delinsValLeuVal*<br>*LeuValAsnThr*<br>*ArgThrSer* | 1 | 57 | 80 | Yes | Yes | - | Crystal<br>structure |

### AlphaFold predictions

Monomer predictions were obtained running AlphaFold v2.0.1 [12, 30], on its simplified version, i.e., with no templates and a reduced Big Fantastic Database (BFD). UCSF ChimeraX v1.3 [31] was used to obtain metrics and figures for AlphaFold results.

We decided to run the aforementioned configuration for AlphaFold as (i) it requires less computational resources; (ii) AlphaFold’s authors claim accuracies to be nearly identical to the regular v2.0.1 for most of the proteins [30], in fact, we see no differences between both configurations (Table 2 and Figure 1); and (iii) antithrombin belongs to the serpin protein superfamily. That means it is not part of the reduced group of proteins showing significant differences between the complete and simplified model, as serpin proteins all have a very similar, conserved structure and are well represented in the databases used to build BFD. Aiming to evaluate the effect of each mutation, we ran an script of Rosetta Online Server that Includes Everyone (ROSIE 2) [32] to stabilize proteins with a point mutation (https://r2.graylab.jhu.edu/apps/submit/stabilize-pm) and we compared the results obtained by both methods. The difference of free energy (ΔΔ*G*) of each protein was calculated in Rosetta Energy Unit (REU). A value of ΔΔ*G* higher than 0 indicates that’s exists a destabilizing mutation.

**Table 2.** Summary of AlphaFold metrics between predictions of reduced and full versions.

| Protein | RMSD mean (Å) | RMSD maximum (Å) | IDDT |
| --- | --- | --- | --- |
| *Wild type* | 0.015 | 0.030 | 0.995 |
| *p.Glu241-Leu242*<br>*delinsValLeuVal*<br>*LeuValAsnThr*<br>*ArgThrSer* | 0.014 | 0.030 | 0.999 |

**Figure 1.**
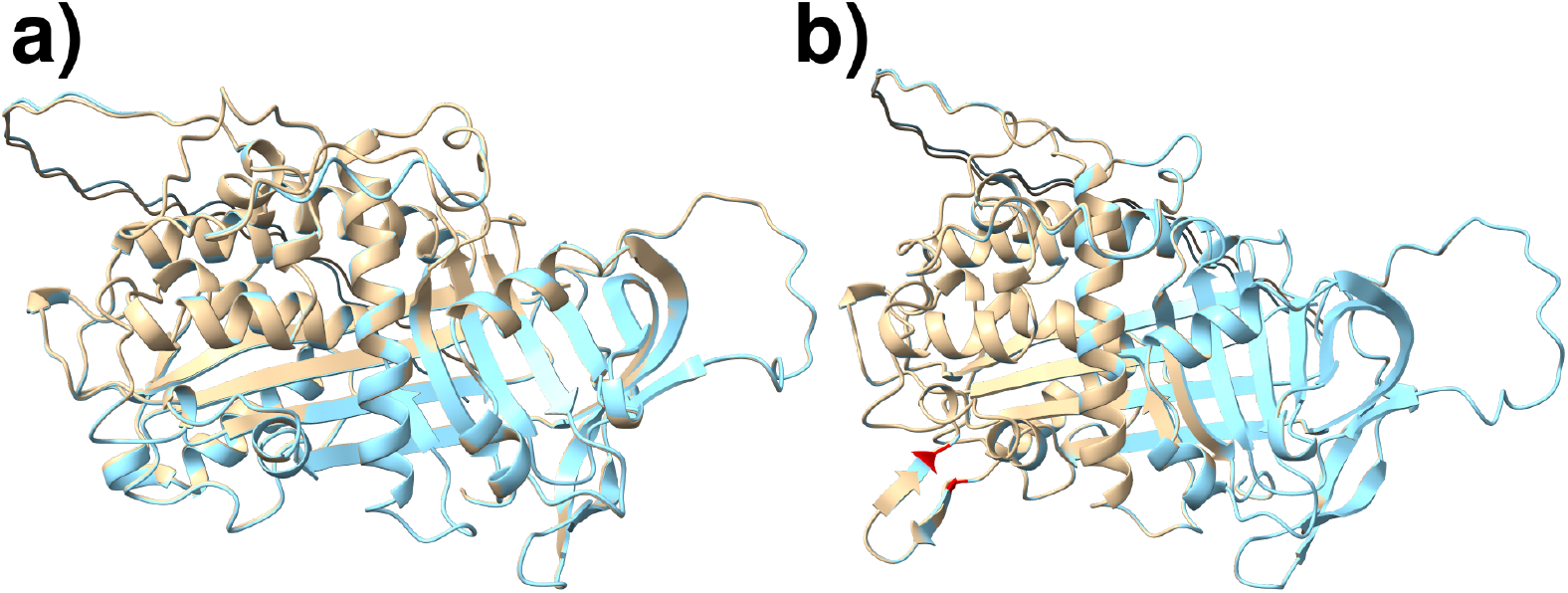
Comparison of AlphaFold’s predictions for wild-type protein (a) and p.Glu241_Leu242delinsValLeuValLeuValAsnThrArgThrSer (b). Reduced version results are colored in grey. Complete version results are in blue. Structural mismatches are colored in red.

### Molecular dynamics

All considered proteins structures were subjected to MD simulations with MD engine Desmond [33] included in the Maestro Suite [34]. All protein models were immersed in a box, whose dimensions where 10×10×10 Å, with water molecules with the simple point charge (SPC) water model. Ions of Sodium and Chlorine were added to neutralize charges and to obtain a molarity of 0.15 M. Energy minimization was made by 2000 steps using the steepest descent method with a threshold of 1.0 kcal/mol/Å. The NPT simulations were realized at the exact temperature in each case: 300, 313 and 373 K with the Nosé-Hoover algorithm [35, 36], and the pressure was maintained at 1 bar with the Martyna-Tobias-Klein barostat [36, 37]. Simulation length of performed MD simulations was 100ns or 1000ns. Periodic conditions were used. The cutoff of 9 Å was established to van der Waals interactions and the Particle Mesh Ewald (PME) method with a tolerance of 10-9 was used in the electrostatic part. The force field used in all runs was OPLS3e [38].

### Metrics

In order to evaluate the accuracy of the structural predictions obtained by AlphaFold and MD simulations, the Root Mean Squared Deviation (RMSD) and the Local Distance Difference Test lDDT [39] metrics were considered for each protein model, when comparing against available crystal structure. In addition, Free Energy Landscape (FEL) analysis combined with PCA and the covariance matrix generation were calculated using GROMACS 2022.2 [40] modules named covar and anaeig when processing Desmond MD trajectories via Maestro Tool: trj convert [11]. Subsequently, 3D FEL figures were obtained using Python library matplotlib 3.5.2 [41]. Pathogenicity predictions were gathered for each variant when available using Varsome v11.3 [42]. Additionally, Ramachandran plots were generated for each structure analyzed as a quality control step (Supplementary Figure 1).

### Code availability

Code used to run AlphaFold is freely available on DeepMind’s GitHub (see references). Software used for MD is property of Schrödinger LCC. Thus, access to it might be restricted. GROMACS and matplotlib codes are available at https://manual.gromacs.org/current/download.html and https://matplotlib.org/stable/users/installing/index.html, respectively.

### Ethical considerations

All methods and experimental protocols used in this study have been carried out in accordance with current guidelines and regulations. This study was approved by the local Ethics Committee of Morales Meseguer University Hospital and performed in accordance with the 1964 Declaration of Helsinki and their later amendments.

All included subjects and/or their legal guardian(s) gave their written informed consent to enter the study.

## Results

### Mutations evaluated

Table 1 shows the mutations selected, the consequences on the protein and the experimental data obtained in each variant.

We selected three missense mutations:

1. p.Arg79Cys. This mutation causes a type II HBS deficiency (Antithrombin Toyama variant) [43], as it changes one arginine residue directly involved in the interaction with heparin [44], the cofactor of antithrombin that fully activates this serpin increasing up to 1000-fold its anticoagulant activity [45]. Moreover, this is one of the most recurrent variants identified in our cohort (N = 19 unrelated cases). All carriers of this variant have reduced heparin cofactor activity, normal antigen levels, and the mutated antithrombin (which constitutes half of the plasma antithrombin) has faster electrophoretic mobility in native-PAGE and low heparin binding under crossed immunoelectrophoresis with heparin. Finally, this residue has also been mutated to Ser or His in other patients with type II HBS deficiency.
2. p.Pro112Ser. This mutation causes a severe reduction of antithrombin levels in plasma, probably by inducing intracellular polymerization according to the detection of disulphide-linked dimers in plasma, as our group previously demonstrated [46].
3. p.Met283Val. The patients carrying this mutation had high antigen levels, with the increase of a form with low heparin affinity, and increased levels of latent antithrombin, all data supporting a type II PE deficiency [25].

We also included in our study two mutations with greater consequences in the amino acid sequence of antithrombin:

1. c.1154-14G>A, that created a new splicing acceptor site in intron 5. The resulting mRNA maintained the reading frame of antithrombin with the insertion of 4 new residues (p.Pro352insValPheLeuPro) [47]. This new variant might induce polymerization of the variant antithrombin, as a severe reduction of antigen levels were observed in 5 unrelated carriers of this recurrent mutation. We also detected the presence of disulphide-linked dimers in plasma, which were slightly bigger than those identified in carriers of the p.Pro112Ser mutation. The purification and proteomic analysis of disulphide-linked dimers from carriers’ plasma confirmed the insertion of the predicted 4 additional amino acids in the variant antithrombin [48].
2. c.722_725delins[731_751;GAACCAG], which caused a complex insertion (p.Glu241_-Leu242delinsValLeuValLeuValAsnThrArgThrSer) in a highly conserved region of serpins. The carrier of this mutation had a type II deficiency with increased levels of a form with low heparin affinity and hyperstable antithrombin. Moreover, structural analysis made with the recombinant protein revealed a relaxed structure with the inserted residues forming a new strand in the central A sheet and a native RCL [49].

### AlphaFold predictions

Firstly, we compared the structure of native antithrombin (PDB: 1AZX) with the one predicted by AlphaFold for the wild type sequence of human antithrombin (https://www.uniprot.org/uniparc/UPI000002C0C1). As shown in Figure 2, the AlphaFold’s predictions were quite accurate, despite some differences concerning the starting or end of helix or strands. Differences are summarized as RMSD in Table 3.

**Figure 2.**
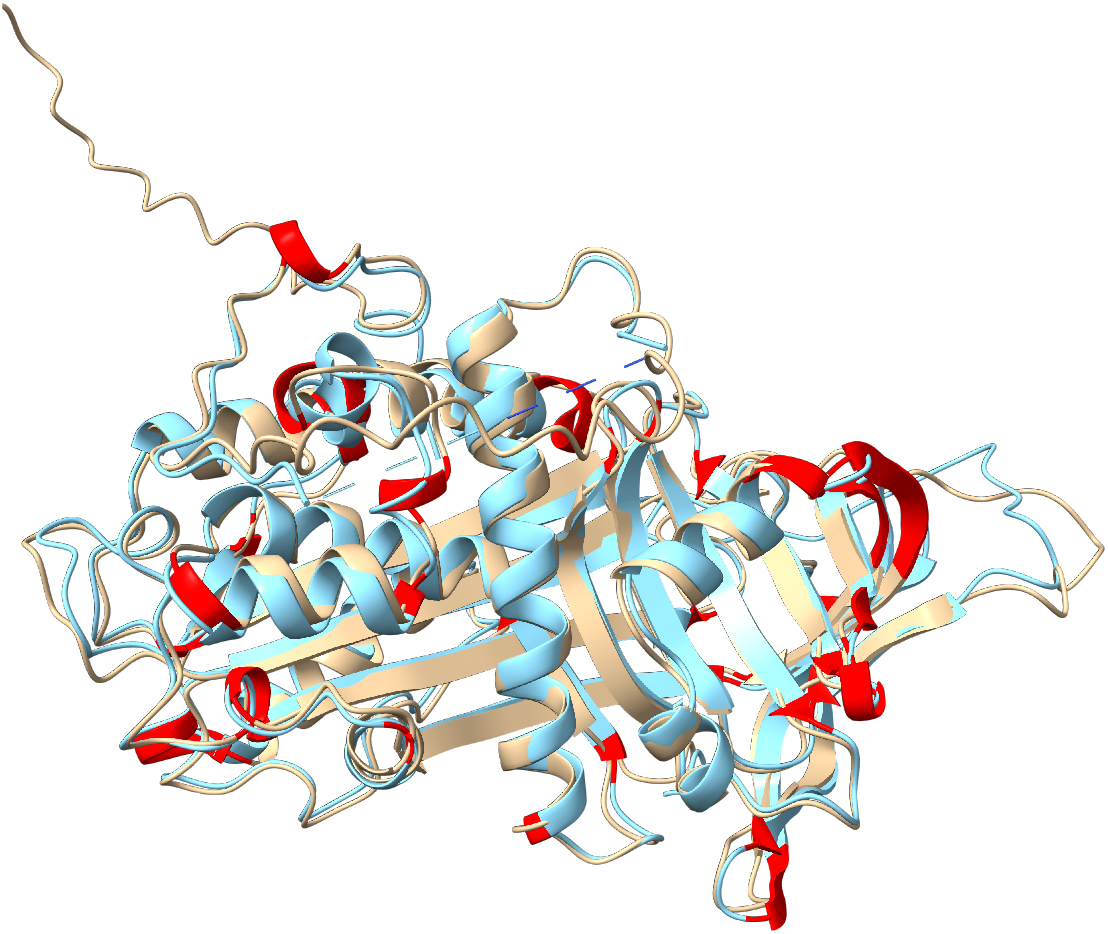
Comparison of crystal structure of wild type native antithrombin (1AZX) (blue) and the structure predicted by AlphaFold for the wild type antithrombin (grey). Mismatches are colored in red.

**Table 3.**
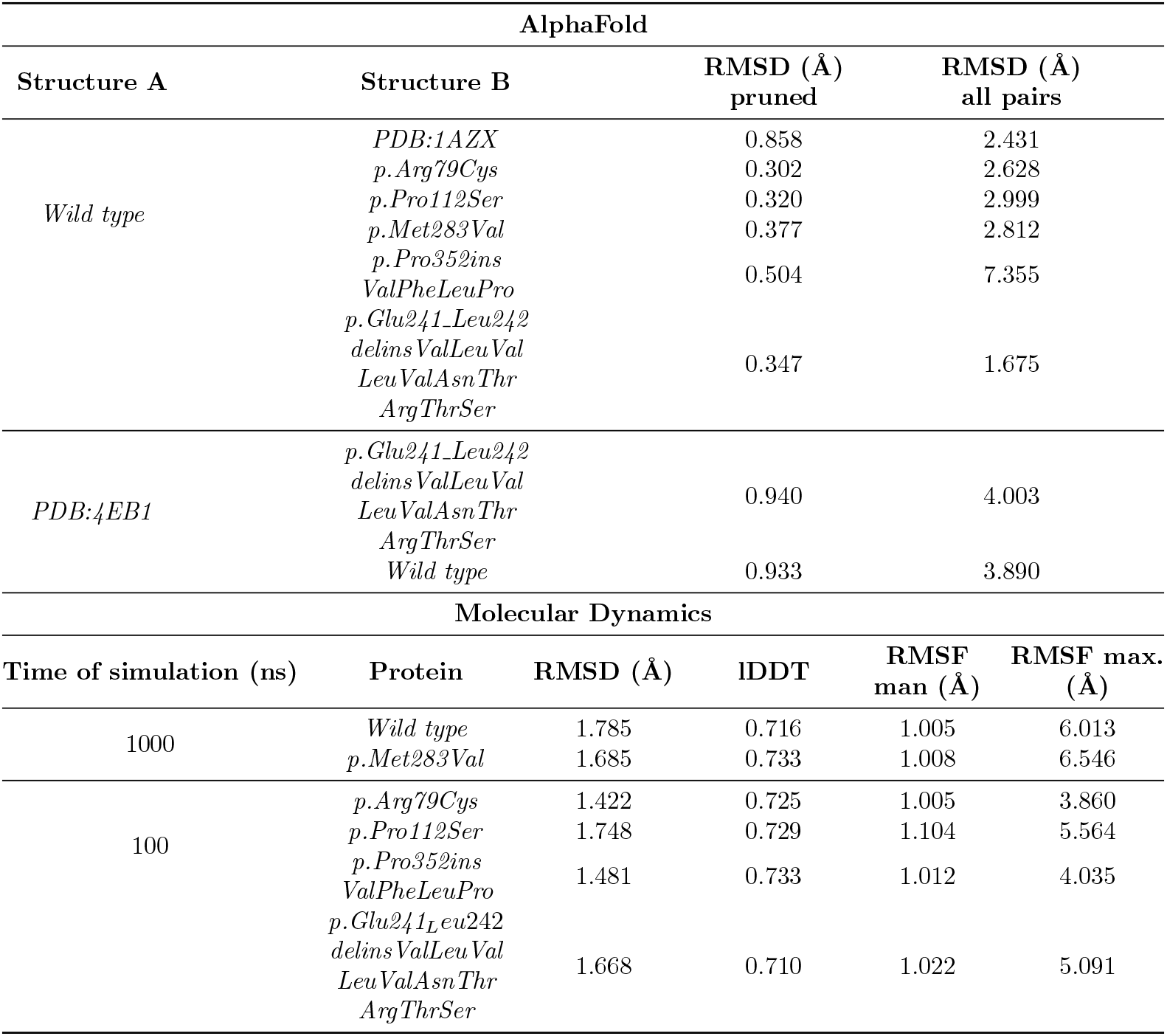
Summary of metrics for AlphaFold and Molecular Dynamics. AlphaFold metrics are referred to its predictions, except for PDB:1AZX and PDB:4EB1. Molecular Dynamics metrics are referred to differences between initial and final state of the proteins.

Then, we compared the AlphaFold’s prediction for the wild type antithrombin with the predictions of the same software for all selected mutants. As expected, the functional variant p.Arg79Cys only caused a minor modification of the structure, even at the mutated residue (Figure 3 and Table 3).

**Figure 3.**
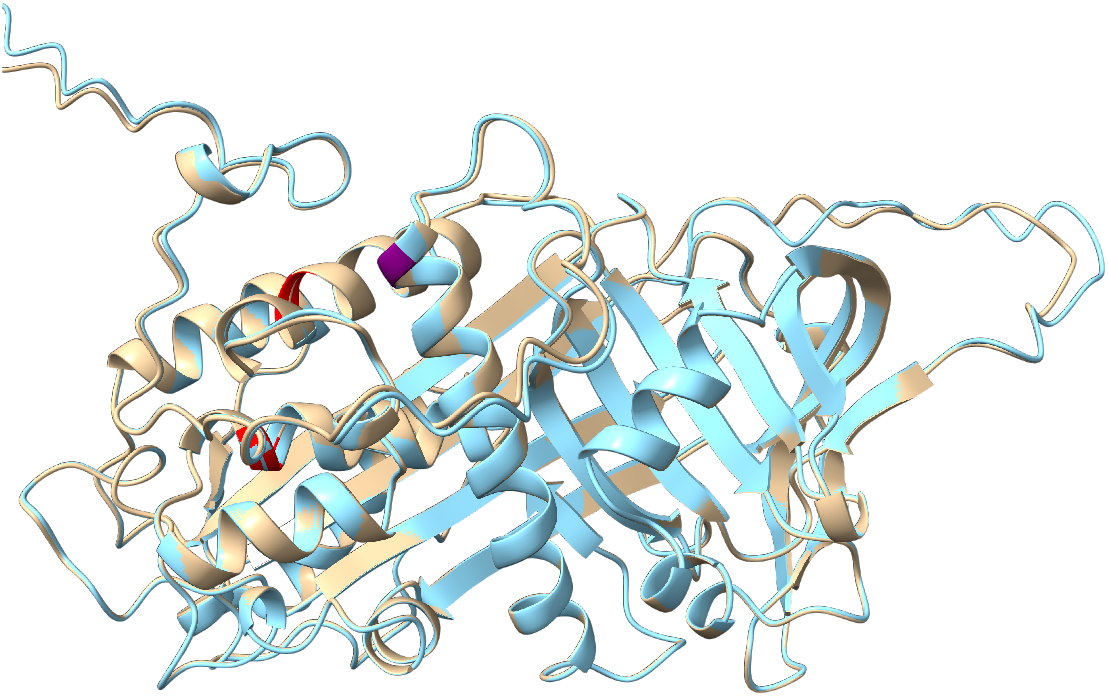
Comparison of AlphaFold predicted structure of wild type antithrombin (grey) and the prediction for p.Arg79Cys, also by AlphaFold (blue). Mismatches are colored in red. The mutated residue is colored in purple.

The comparison of the other two missense mutations included in this study was quite similar, with minor mismatches (Figure 4 and Table 3). These three mutations (p.Arg79Cys, p.Pro112Ser and p.Met283Val) were analyzed using Rosetta Online Server that Includes Everyone (ROSIE 2). The results showed that all mutations have a value of REU higher than the native (Table 4), thus all were destabilizing, specially p.Pro112Ser, with an energy difference of 5.88 REUp.Arg79Cys and p.Met283Vals have an energy difference close to 2 REU. Furthermore, we compared ROSIE 2 predictions with AlphaFold’s. These comparisons are summarized in Table 4.

**Figure 4.**
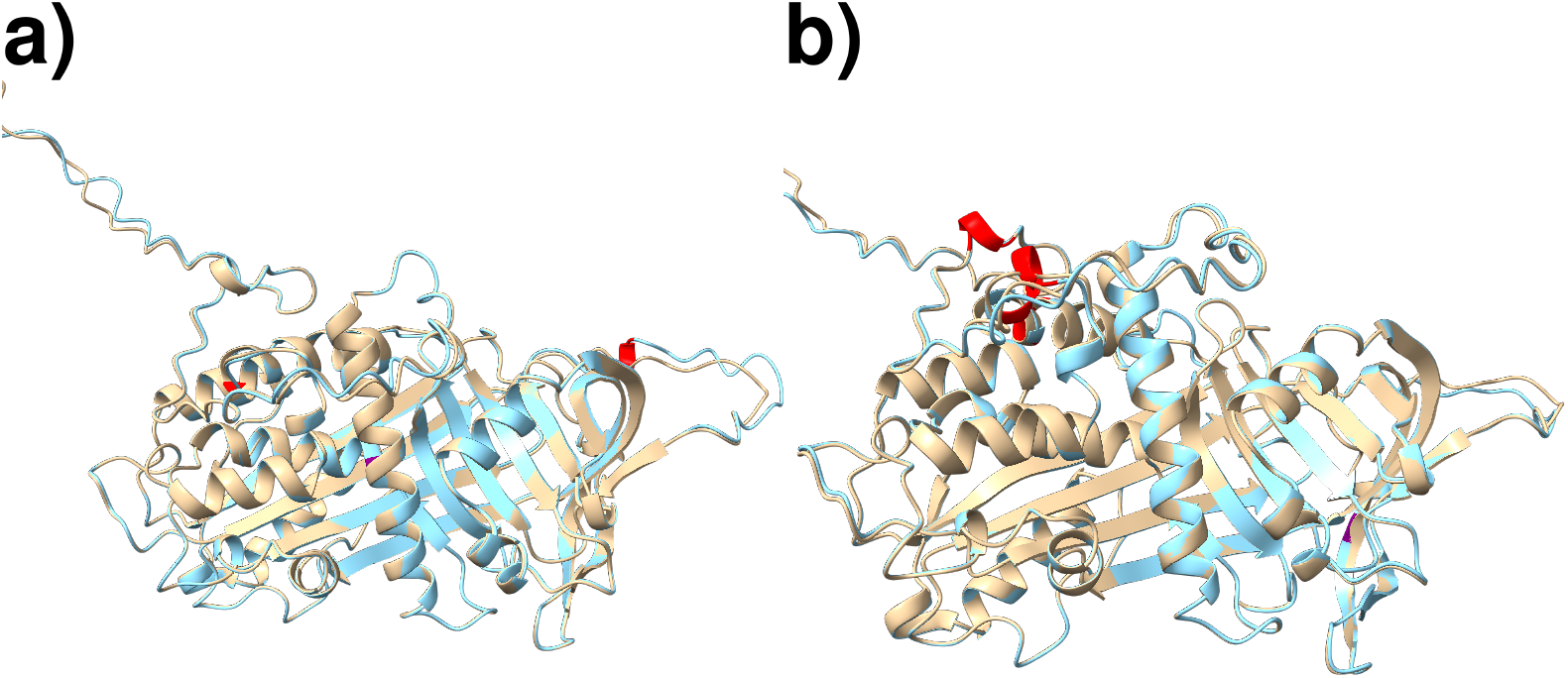
Comparison of AlphaFold predicted structure of wild type antithrombin (grey) and the prediction for p.Proll2Ser (a) and p.Met283Val (b), also by AlphaFold (blue). Mismatches are colored in red. The mutated residue is colored in purple.

**Table 4.**
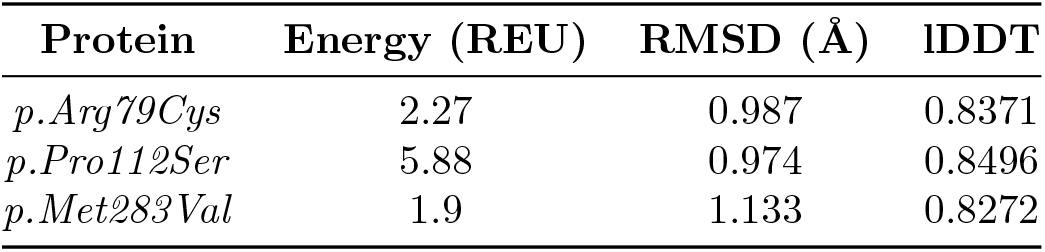
Summary of RMSD metrics of the difference of energy in REU (Rosetta Energy Units) between the wild type and each mutant. Differences between ROSIE 2 predictions and AlphaFold’s are referred as RMSD and lDDT.

Interestingly, AlphaFold also produced a native structure for variants causing insertions of residues. For the small insertion of 4 amino acids caused by the intronic mutation (p.Pro352insValPheLeuPro), AlphaFold predicted a small elongation of the loop connecting hI and s5A (Figure 5 and Table 3).

**Figure 5.**
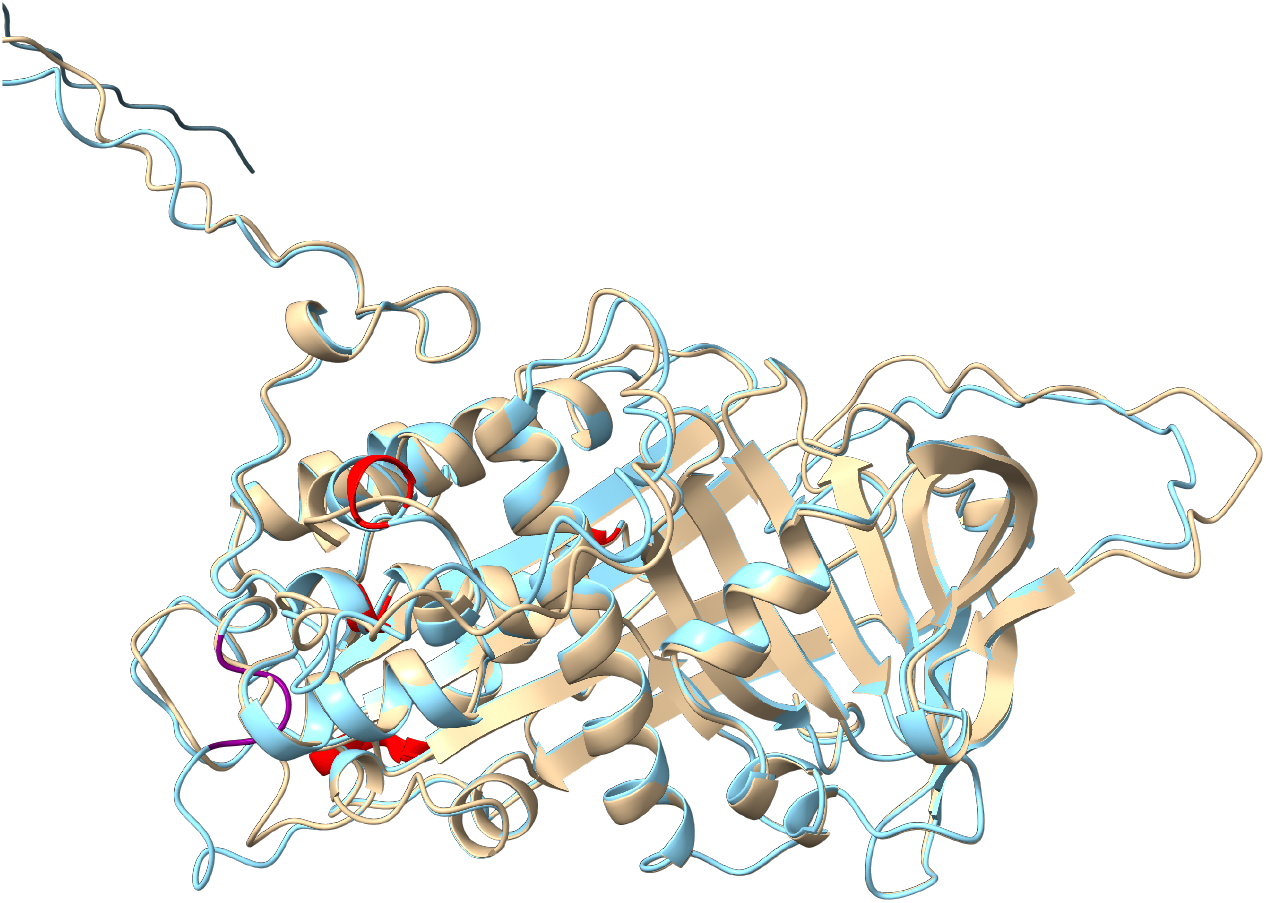
Comparison of AlphaFold predicted structure of wild type antithrombin (grey) and the prediction for p.Pro352insValPheLeuPro, also by AlphaFold (blue). Mismatches are colored in red. The mutated residues are colored in purple.

For the p.Glu241_Leu242delinsValLeuValLeuValAsnThrArgThrSer variant, the inserted new residues are forced into a new small helix but maintaining the stressed native structure, which differed significantly from the crystal structure of this variant (4EB1) (Figure 6 and Table 3).

**Figure 6.**
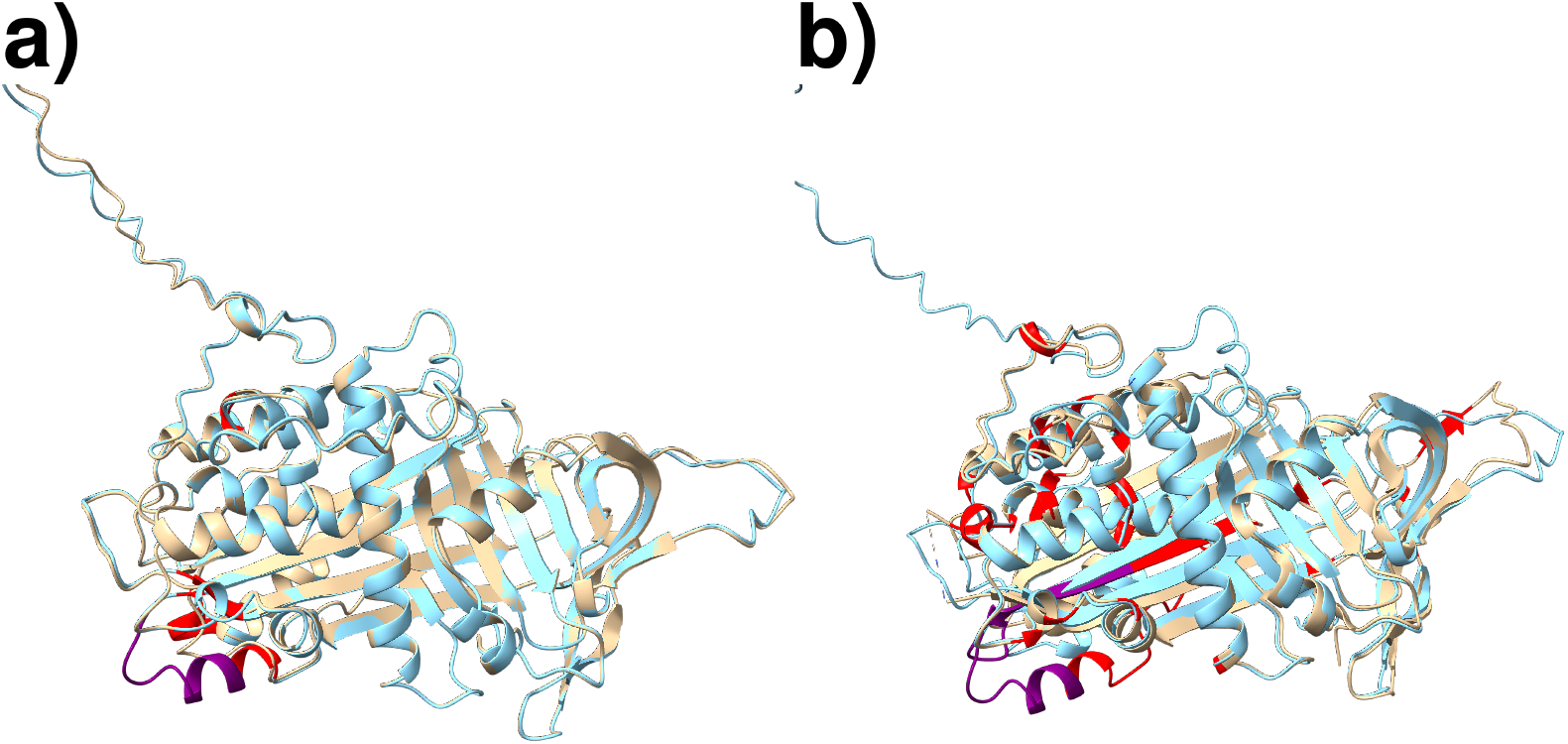
Comparison of AlphaFold prediction for p.Glu241_Leu242delinsValLeuValLeuValAsnThrArgThrSer (blue) with the wild type antithrombin (grey) (a) and the crystal structure of this variant (4EB1, in grey) (b). Mismatches are colored in red. The mutated residues are colored in purple.

When analyzing changes in the hydrogen bond network of the different structures, we observed little differences between the hydrogen network of AlphaFold’s wild-type and its predictions for point mutations (Supplementary Figure 2e-g), sharing near 90% of intra-chain connections via hydrogen bonds. Information is summarized in Figure 7a and 7b.

**Figure 7.**
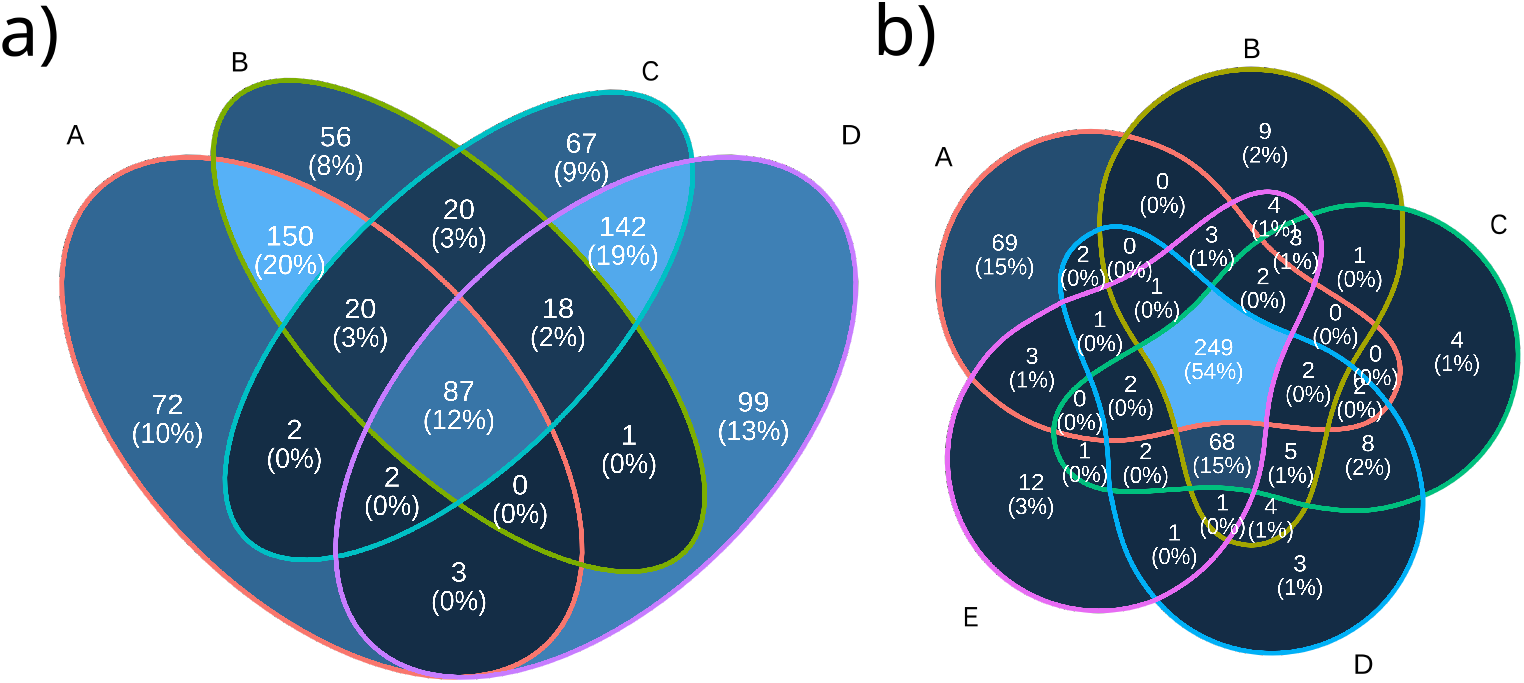
Analysis of the hydrogen bond network. Comparison of pairs of residues linked by at least one hydrogen bond in each structure. a) Changes between wild-type and p. Glu241_Leu242delinsValLeuValLeuValAsnThrArgThrSer: A) 1AZX-I, B) AlphaFold’s wild-type, C) AlphaFold’s p. Glu241_Leu242delinsValLeuValLeuValAsnThrArgThrSer, D) 4EB1-I. b) Changes between wild-type and point mutations: A) 1AZX-I, B) AlphaFold’s wild-type, C) AlphaFold’s p.Arg79Cys, D) AlphaFold’s p.Pro112Ser, E) AlphaFold’s p.Met283Val

Nonetheless, when comparing 1AZX’s hydrogen bond network with 4EB1’s (Supplementary Figure 2b) and AlphaFold’s p. Glu241_Leu242delinsValLeuValLeuValAsnThr-ArgThrSer (Supplementary Figure 2c) we find the latter having more in common with the native crystal than 4EB1.

Also, we evaluated the differences between the reduced and complete version of AlphaFold with both wild-type protein and p.Glu241_Leu242delinsValLeuValLeuValAsn-ThrArgThrSer. We selected the wild-type sequence for benchmarking as AlphaFold has been validated with wild-type proteins, and p.Glu241_Leu242delinsValLeuValLeuValAsn-ThrArgThrSer as it is the most aberrant and challenging variant for modelling of our study. Resulting metrics (Table 2) showed virtually nonexistent differences between both versions of AlphaFold, validating thus the applicability of the reduced version of AlphaFold, as its results are comparable to those obtained with the complete version of the model.

### Molecular dynamics

The overall structure of wild type antithrombin did not change significantly with simulations at 1000 ns and low (27ºC), high (40ºC) or extreme temperatures (100ºC), with the RCL released, showing a stressed structure (Figure 8). Indeed, the structure was also similar (stressed native) for variants secreted with a relaxed structure (p.Met283Val and p.Glu241_Leu242delinsValLeuValLeuValAsnThrArgThrSer) with minor changes near the mutations (Figure 8b and 8c). Metrics are summarized in Tables 3–6.

**Figure 8.**
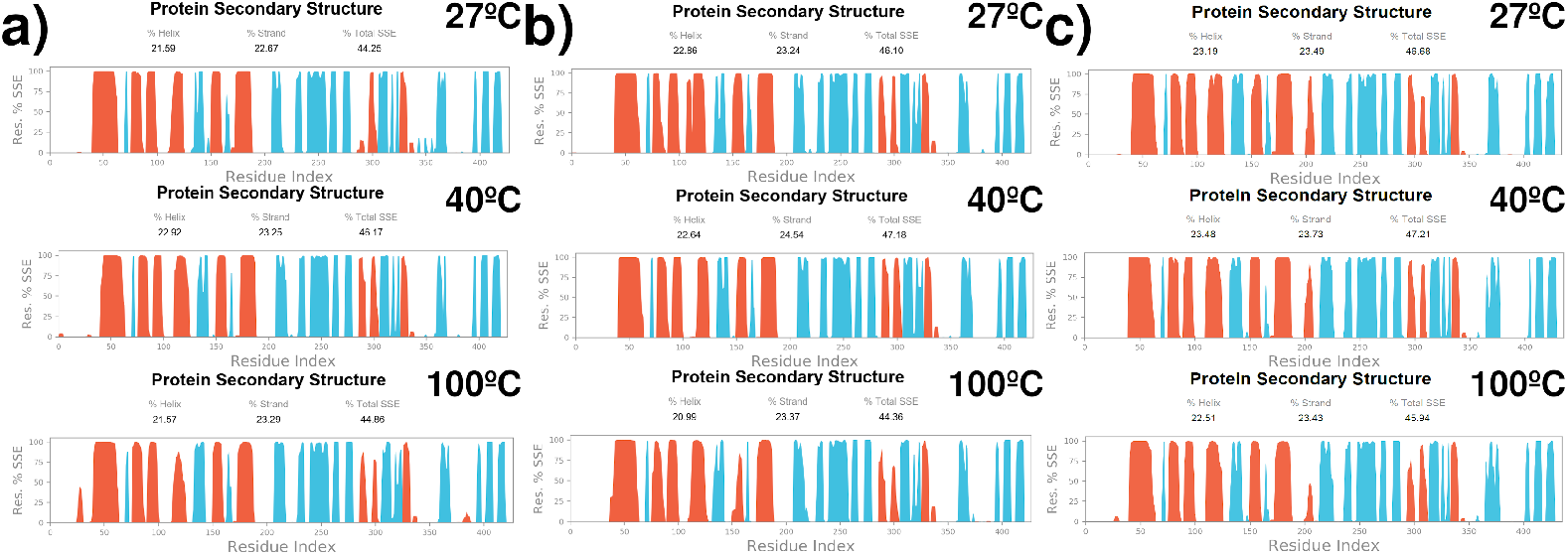
Molecular dynamics predictions obtained during 1000 ns for wild type antithrombin (a) and conformational variants secreted in a relaxed conformation, p.Met283Val (b) and p.Glu241_Leu242delinsValLeuValLeuValAsnThrArgThrSer (c) at 27, 40 and 100ºC.

**Table 5.**
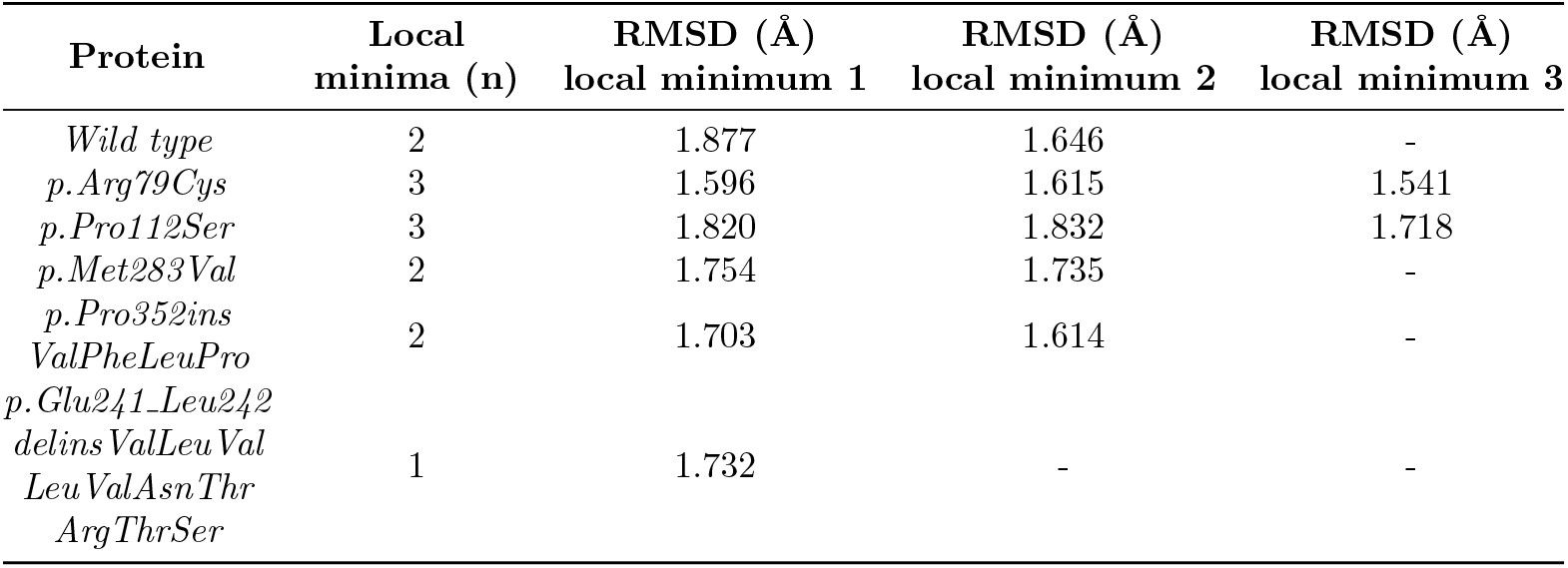
Summary of RMSD metrics between initial and most energetically stable MD states for all studied proteins. Local minimums were calculated using FEL calculations.

**Table 6.**
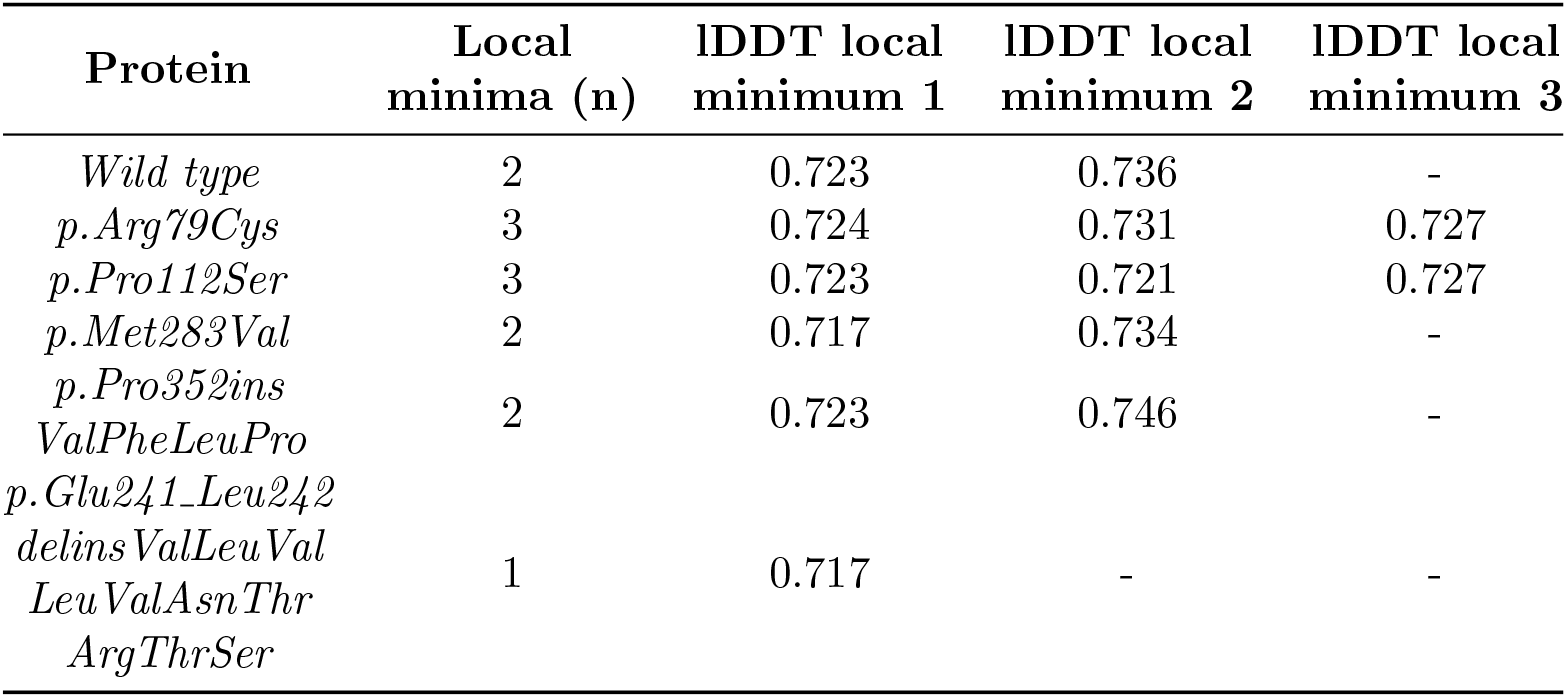
Summary of lDDT metrics between initial and state with less energy for all considered proteins.

We also studied the different proteins using the FEL approach [50] to find the most energetically stable pose within the trajectory, and comparing its conformation against the initial from MD trajectory. These conformations are local minimums. A summary of obtained results is shown in Tables 5–6. Details regarding 3D FEL figures are presented in Supplementary Figure 3.

## Discussion

The structural flexibility of serpins, required for their efficient inhibitory mechanism, also make these molecules particularly sensitive to environmental conditions (pH, temperature, redox conditions) with both physiological and pathological consequences [51, 52, 53, 54, 55, 56]. Even minor missense mutations might disturb the stressed native conformation and ease the transition to relaxed, hyperstable non-inhibitory conformations with the RCL inserted (mainly latent or polymers) [3]. However, there are other mutations that, affecting RNA stability or translation cause quantitative deficiency of serpins and could have pathological consequences. Finally, there are also mutations affecting functional domains that, with minor effects in folding or secretion, cause a qualitative deficiency with inactive variants also involved in different disorders [8]. Hemostatic serpins, particularly antithrombin, are excellent examples of the pathogenic impact of these environmental conditions and types of mutations [3, 9, 57]. Thus, it would be of great interest to have tools able to predict the effect of mutations affecting serpins, particularly those causing transitions to the latent or polymer conformations as they seem to have higher clinical severity [58], probably by a dominant negative effect additional to the loss of function associated to these mutations [59, 60], and by potential additional deleterious consequences if accumulated intracellularly.

In this study, we explored the predictions of AlphaFold, a novel artificial intelligence system conceived to predict structures of new proteins, and molecular dynamics, a computer simulation method for analyzing the physical movements of atoms and molecules on the consequences of five *SERPINC1* mutations causing antithrombin variants, four of them with experimental evidences supporting a conformational change, as well as the effect of high temperatures that caused conformational changes even in wild type antithrombin [51].

AlphaFold gave an accurate prediction of the wild type antithrombin structure. Similarly, p.Arg79Cys, a recurrent functional mutation that impairs the binding of antithrombin to its cofactor heparin, caused a minor defect in the structure predicted by AlphaFold. However, AlphaFold also maintained the native conformation for all other mutations with experimental evidence supporting a strong conformational consequence, both polymer/dimer formation (p.Pro112Ser, and p.Pro352insValPheLeuPro), latent transition (p.Met283Val) or a new relaxed structure formed as consequence of a relatively large insertion (p.Glu241_Leu242delinsValLeuValLeuValAsnThrArgThrSer). It has been argued that AlphaFold has not been designed to predict the effect of SNVs [19], but as shown for the cases with 4 to 10 residues insertion, using either reduced or full versions, it only adapts the inserted residues to the structure with minor deviations of the native conformation. Thus, for the in-frame insertion of 4 residues (p.Pro352insValPheLeuPro) AlphaFold expands a loop to include the new 4 residues, and for the more complex variant p.Glu241_Leu242delinsValLeuValLeuValAsnThrArgThrSer, a small helix containing the 10 inserted residues was generated.

Molecular dynamics also maintain the native conformation for all variants causing relaxed structures, even creating a new helix for the more complex variant, like AlphaFold does. But the prediction also yielded the native conformation when the experiments were executed under environmental conditions that exacerbate or directly induced conformational changes of mutants and even the wild type molecule (40ºC and 100ºC).

Thus, these two predictive tools forced the folding of antithrombin to the native stressed conformation even for mutations or conditions that render relaxed structures with experimental evidence.

The explanation to these incorrect predictions may be found in the PDB database. This database contains 334 entities annotated as serpins (with PFAM identifier PF00079), most of them in the native conformation. Indeed, only 36 of those remain when searching for latent. Interestingly, for human antithrombin, the first crystal structure obtained simultaneously by two independent groups, contains a dimer of a latent and a native molecule [61, 62, 63]. This misbalance of structures with native conformation, and the strong effect of using a specific structure for molecular dynamics studies may lead these predictive tools to force a native structure as the final result of all predictions, even over the own crystal as it occurs for the p.Glu241_Leu242delinsValLeuValLeuValAsnThrArg-ThrSer complex insertion [49]. Therefore, it is necessary to improve predictive tools of folding for serpins, which must consider the conformational sensitivity of these molecules and the transition to hyperstable conformations as the most feasible folding associated to disturbing mutations. In the case of MD simulations, we hypothesize these limitations might be overcome if we could reach longer time scales where such structural rearrangements occur, using for that specialized and private supercomputers such as ANTON-3 [64], or applying special MD techniques that force sampling of specific events such as accelerated MD or Metadynamics [65, 66, 67]. Nonetheless, MD results of all proteins showed no significant differences. If we could run multiple MD with different initial seeds, perhaps we could appreciate changes between MD for each protein. Furthermore, recent implementations of novel machine learning methods coupled with molecular dynamics may improve the research on the proteins’ conformational ensemble [11]. As for AlphaFold, an in-depth study of the method and understanding of the code and training process might allow providing protein-targeted predictions for this family of serpin structures with higher accuracy [68].

In this work we assessed whether AlphaFold could generalize outside of its established range of applicability. It is important to clearly define the capabilities and limitations of AlphaFold, given that if the model was able to generate 3D structures of mutated proteins it would be a huge step forward in several areas of Biomedical research. Rare Diseases research is an example, where given the limited number of patients and resources, it is not always feasible to crystallize mutant proteins in order to assess the pathogenicity of observed variants. In such cases, having access to a model capable of predicting the structural consequences of mutations would be a great improvement.

Despite other studies have analyzed AlphaFold capabilities with point, simulated mutations, this study includes actual variants, found in patients. Furthermore, we do not only include point mutations, like previous studies have done [19]. Instead, we also include bigger rearrangements (it is the case of variant p.Glu241_Leu242delinsValLeuValLeuValAsn-ThrArgThrSer, of which we also have access to its crystal, 4EB1 [49]) to assess AlphaFold’s capability to generalize out of the serpin structures the model has already seen on its training.

Moreover, AlphaFold is partially based on a neural network. Such systems are a *black box model*, making it difficult to guess how they take decisions or why they create a certain output. Thus, running it with different inputs (in this case, sequences of different nature, wild-type, point mutations, rearrangements) may give some clues of how the network is internally working. The results of variants p.Pro352insValPheLeuPro and p.Glu241_Leu242delinsValLeuValLeuValAsnThrArgThrSer suggest that AlphaFold is creating relatively simple structures as loops or small helices to accommodate the mutated sequence outside of the main structure, so it does not obstruct in the reconstruction of the native templates seen during the model’s training. In summary, we observed that, contrary to what might appear looking at the outstanding results of AlphaFold in the two last CASPs with wild-type proteins [14, 16], the old protein folding problem remains, in some respects, unsolved.

## Acknowledgements

The study was supported by grants from Instituto de Salud Carlos III, FEDER (PI21/00174 & PMP21/00052) and Fundación Séneca de la Región de Murcia under Project 20988/PI/18; Garrido-Rodríguez P and de la Morena-Barrio B hold contracts from CIBERER. Carmena-Barguenño M is a predoctoral student employed to the training research staff financed by Plan Propio de Investigación de la UCAM. Bravo-Pérez C has a Río Hortega fellowship (CM20/00094). Cifuentes-Riquelme R has a PFIS contract. de la Morena-Barrio ME has a postdoctoral contract from University of Murcia. This research was partially supported by the supercomputing infrastructure of the NLHPC (ECM-02), Powered@NLHPC, by the Plataforma Andaluza de Bioinformática of the University of Málaga, and the Extremadura Research Centre for Advanced Technologies (CETA-CIEMAT), funded by the European Regional Development Fund (ERDF). CETA-CIEMAT is part of CIEMAT and the Government of Spain.

## Author contributions

AlphaFold was run by PGR. Molecular Dynamics simulations were performed by MCB and HPS. Selection of variants was done by MEMB, JC and CBP. Cohort managing and wet-lab assays were performed by MEMB, BMB, CBP, RCR, MLL and JC. Pathogenicity predictions were gathered by MEMB. Downstream analysis of results was performed by PGR. Manuscript was written by PGR, MCB, HPS and JC. All authors revised the document and gave their feedback.

## Data availability

Variants p.Arg79Cys, p.Pro112Ser, p.Met283Val, p.Pro352insValPheLeuPro and p.Glu-241_Leu242delinsValLeuValLeuValAsnThrArgThrSer, mentioned in our work, are deposited in UniProt as VAR_007037, VAR_086227, VAR_027468, VAR_086198 and VAR_086197, respectively.

## Additional information

The authors declare no competing interests.

