## Supplementary Figure 1 for "Analysis of AlphaFold and molecular dynamics structure predictions of mutations in serpins"

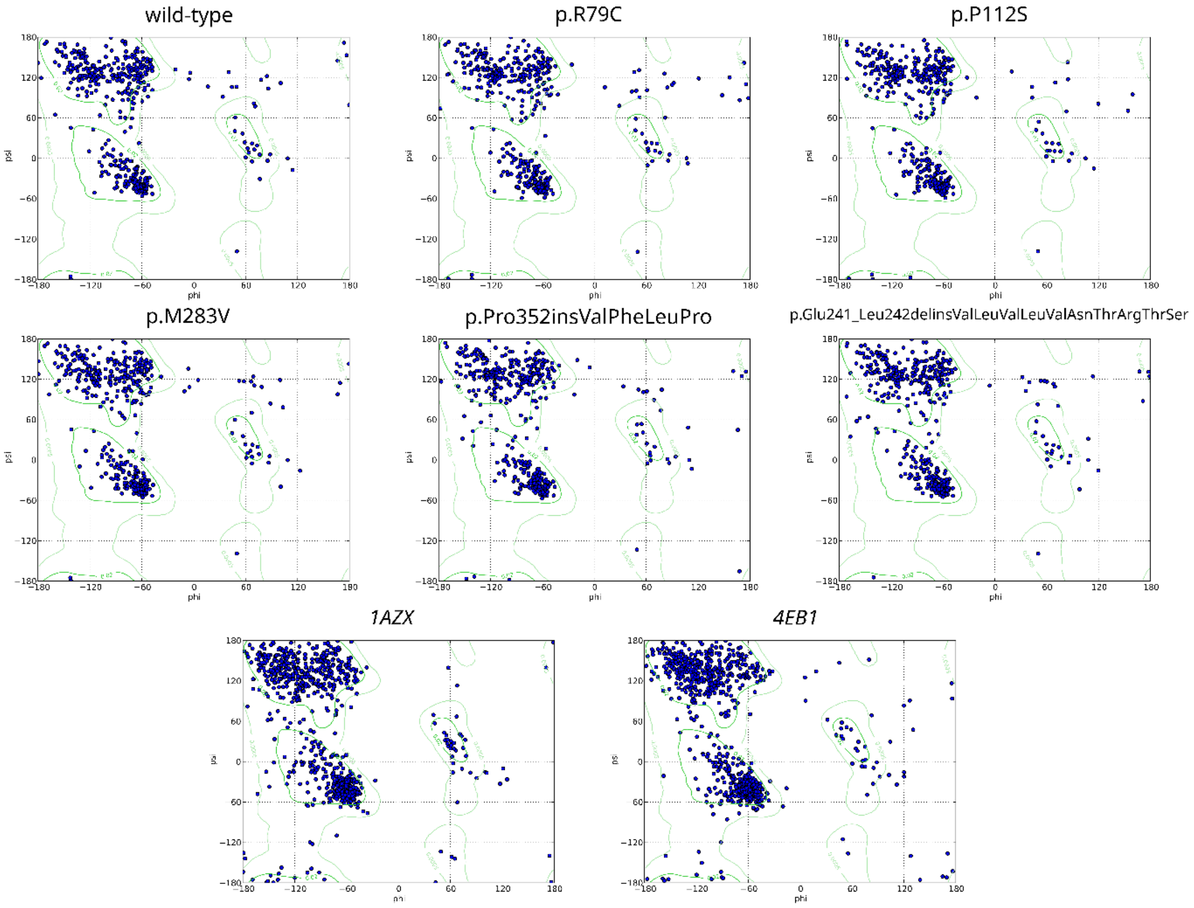


**Supplementary Figure 1.** *Ramachandran plots for structures analyzed. Mutant structures correspond to AlphaFold's prediction for said variant. Plots entitled in caps and italics refer to stated PDB crystal structures.*
