## Supplementary Figure 2 for "Analysis of AlphaFold and molecular dynamics structure predictions of mutations in serpins"

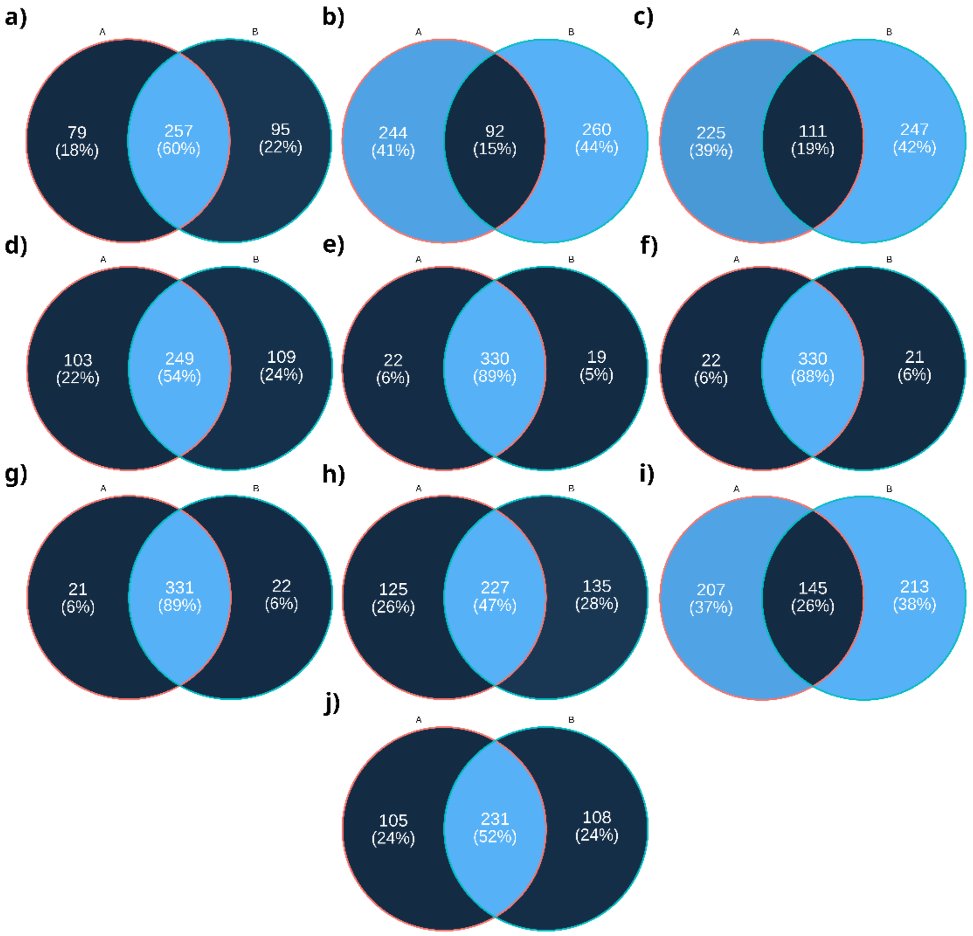


**Supplementary Figure 2.** *Binary comparisons of pairs of residues linked by, at least, one hydrogen bond. Non-crystal structures refer to AlphaFold predictions. a) 1AZX-I vs wild-type, b) 1AZX-I vs 4EB1-I, c) 1AZX-I vs p. Glu241_Leu242delinsValLeuValLeuValAsnThrArgThrSer, d) 4EB1-I vs p. Glu241_Leu242delinsValLeuValLeuValAsnThrArgThrSer, e) wild-type vs p.Arg79Cys, f) wild-type vs p.Pro112Ser, g) wild-type vs p.Met283Val, h) wild-type vs p. Pro352insValPheLeuPro, i) wild-type vs p. Glu241_Leu242delinsValLeuValLeuValAsnThrArgThrSer, j) 1AZX-I vs 1AZX-L.*
