## Supplementary Figure 3 for "Analysis of AlphaFold and molecular dynamics structure predictions of mutations in serpins"

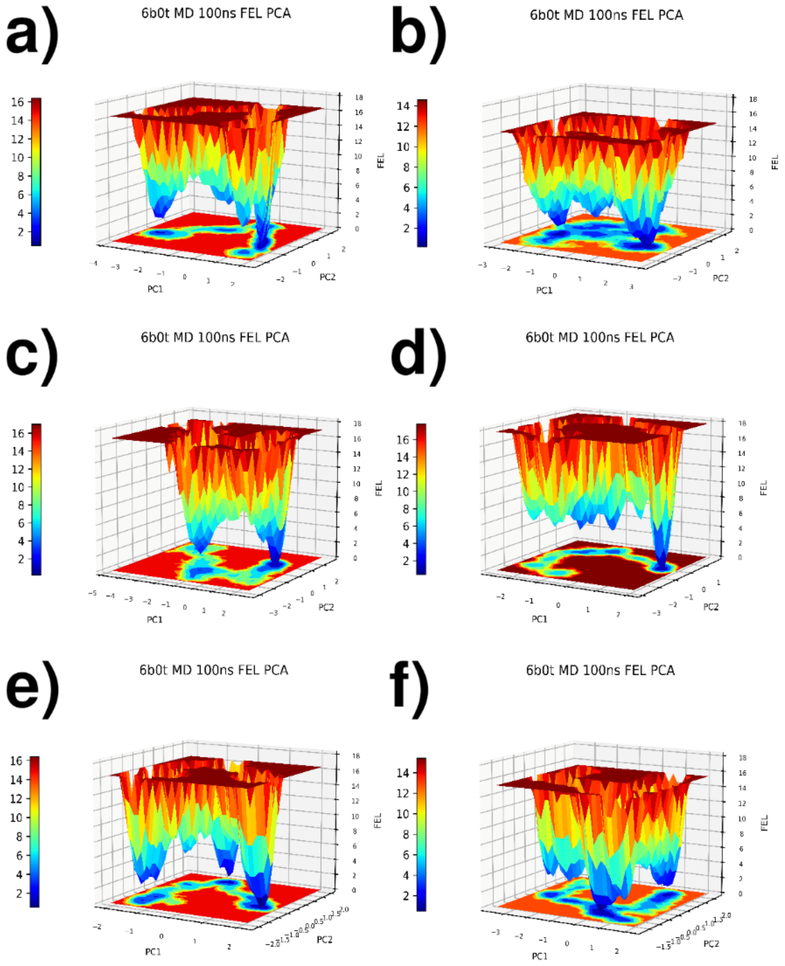


**Supplementary Figure 3.** Free Energy Landscape profiles for all studied proteins after analysis of their MD trajectories. X and Y axis represent PC1 and PC2 PCA components and Z axis free energy value. a) Wild type, b) p.Arg79Cys, c) p.Pro112Ser, d) p.Met283Val, e) p. Pro352insValPheLeuPro, f) p. Glu241_Leu242delinsValLeuValLeuValAsnThrArgThrSer.
