## Supplementary Table 1 for "Analysis of AlphaFold and molecular dynamics structure predictions of mutations in serpins"

| Variant | Origin | Prediction | Score |
|---|---|---|---|
| p.Arg79Cys | ACMG | Pathogenic | - |
| | MetaLR | Damaging | 0.6737 |
| | MetaSVM | Damaging | 0.4294 |
| | MetaRNN | Damaging | 0.8396 |
| | REVEL | Pathogenic | 0.7429 |
| | BayesDel addAF | Damaging | 0.1701 |
| | BayesDelnoAF | Damaging | 0.2987 |
| | DEOGEN2 | Damaging | 0.8176 |
| | EIGEN | Pathogenic | 0.6868 |
| | EIGEN PC | Pathogenic | 0.6482 |
| | FATHMM | Damaging | -1.69 |
| | FATHMM-MKL | Damaging | 0.9084 |
| | FATHMM-XF | Damaging | 0.9012 |
| | LIST-S2 | Damaging | 0.9565 |
| | LRT | Deleterious | 0 |
| | M-CAP | Damaging | 0.1211 |
| | MVP | Pathogenic | 0.9571 |
| | MutPred | Pathogenic | 0.755 |
| | Mutation assessor | Medium | 2.19 |
| | MutationTaster | Disease causing | 1 |
| | PROVEAN | Damaging | -2.68 |
| | PrimateAI | Tolerated | 0.7381 |
| | SIFT | Damaging | 0 |
| | SIFT4G | Damaging | 0.001, 0.002 |
| p.Pro112Ser | ACMG | Pathogenic | - |
| | MetaLR | Damaging | 0.9576 |
| | MetaSVM | Damaging | 10.973 |
| | MetaRNN | Damaging | 0.9775 |
| | REVEL | Pathogenic | 0.9649 |
| | BayesDel addAF | Damaging | 0.5951 |
| | BayesDel noAF | Damaging | 0.617 |
| | DEOGEN2 | Damaging | 0.9388 |
| | EIGEN | Pathogenic | 0.9768 |
| | EIGEN PC | Pathogenic | 0.9297 |
| | FATHMM | Damaging | -4.28 |
| | FATHMM-MKL | Damaging | 0.9387 |
| | FATHMM-XF | Damaging | 0.9657 |
| | LIST-S2 | Damaging | 0.9547, 0.9508 |
| | LRT | Deleterious | 0 |
| | M-CAP | Damaging | 0.3203 |
| | MVP | Pathogenic | 0.9766 |
| | MutPred | Pathogenic | 0.932 |
| | Mutation assessor | Medium | 3.09 |
| | MutationTaster | Disease causing | 1 |
| | PROVEAN | Damaging | -6.64 |
| | PrimateAI | Tolerated | 0.7232 |
| | SIFT | Damaging | 0 |
| | SIFT4G | Damaging | 0, 0.019 |
| p.Met283Val | ACMG | Uncertain significance | - |

| | | | |
|---|---|---|---|
| | MetaLR | Damaging | 0.6121 |
| | MetaSVM | Damaging | 0.1556 |
| | MetaRNN | Damaging | 0.8389 |
| | REVEL | Pathogenic | 0.689 |
| | BayesDel addAF | Damaging | 0.2569 |
| | BayesDel noAF | Damaging | 0.1314 |
| | DEOGEN2 | Damaging | 0.9168 |
| | EIGEN | Benign | -0.0229 |
| | EIGEN PC | Benign | 0.006645 |
| | FATHMM | Damaging | -2.37 |
| | FATHMM-MKL | Damaging | 0.9347 |
| | FATHMM-XF | Damaging | 0.8248 |
| | LIST-S2 | Tolerated | 0.7519 |
| | LRT | Deleterious | 0 |
| | M-CAP | Damaging | 0.07539 |
| | MVP | Pathogenic | 0.9613 |
| | MutPred | Pathogenic | 0.942 |
| | Mutation assessor | Medium | 3.24 |
| | MutationTaster | Disease causing | 0.9997 |
| | PROVEAN | Damaging | -2.77 |
| | PrimateAI | Tolerated | 0.3828 |
| | SIFT | Damaging | 0.019 |
| | SIFT4G | Damaging | 0.037 |
| | EVE | Uncertain | 0.4142 |
| p.Pro352insValPheLeuPro | ClinVar | Pathogenic | - |
| p.Glu241_Leu242delinsValLeuValLeuValAsnThrArgThrSer | - | - | - |

**Supplementary Table 1.** Pathogenicity predictions for selected variants.
